# Semaglutide drives weight loss through cAMP-dependent mechanisms in GLP1R-expressing hindbrain neurons

**DOI:** 10.1101/2025.08.12.668772

**Authors:** Claire Gao, Isabelle C. Geneve, Chia Li, Kaitlyn McElhern, Marc L. Reitman, Andrew Lutas, Michael J. Krashes

## Abstract

Glucagon-like peptide 1 receptor (GLP1R) agonists like semaglutide drive weight loss through the brain, but insights into their intracellular signaling mechanisms are lacking. Although canonically GLP1Rs signal through the stimulatory alpha Gs protein, we find that semaglutide utilizes both Gs- and Gq-signaling pathways in GLP1Rs in the area postrema (AP)—the primary site of semaglutide action in the brain—to differentially regulate neuronal activation across distinct neuronal clusters. Semaglutide also drives graded increases of the essential secondary messenger cyclic adenosine monophosphate (cAMP) in Glp1r-expressing AP neurons (AP^Glp1r^) through Gs-dependent and -independent manners. Inhibition of the cAMP-degradation enzyme phosphodiesterase 4 (PDE4) can enhance and sustain these cAMP responses, whereas disruption of Gs or cAMP signaling in AP^Glp1r^ neurons abolishes semaglutide-induced weight loss and downstream brain-wide activation. Our systematic characterization of semaglutide’s signaling mechanisms in the brain provides avenues for improving the performance of obesity therapeutics.

## Introduction

Obesity is a complex and costly epidemic with increasing international prevalence^1–3^. As such, there is high demand for a family of anti-obesity drugs known as glucagon-like peptide 1 receptor (GLP1R) agonists (GLP1RAs) such as semaglutide, which effectively reduce body weight and food intake in rodent models and humans^4,5^. Despite an emerging consensus that these peripherally administered GLP1RAs exert their effects on appetite suppression through the central nervous system, knowledge of the neural pathways and intracellular signaling mechanisms by which they act in the brain remains limited.

Studies in mice showed that pharmacological blockade or genetic knockout of GLP1Rs in the central nervous system (CNS) could block or significantly attenuate GLP1RA weight loss and food intake suppression, highlighting the necessity of central GLP1Rs^6–9^. Systemic GLP1RA injections lead to a distributed network of neuronal activation of regions—such as parabrachial nucleus (PBN), central amygdala (CeA), area postrema (AP), nucleus of the solitary tract (NTS), bed nucleus of the stria terminalis (BNST)—implicated in central appetite regulation^10–13^. However, systemically injected labeled GLP1RAs show limited brain penetration, with the strongest labeling observed in circumventricular organs such as the AP and arcuate nucleus (ARC), suggesting that most central effects are mediated by downstream signaling from these primary GLP1RA binding sites^14^. Among these, Glp1r-expressing neurons in the hindbrain, particularly in the dorsal vagal complex composed of AP and NTS, have been identified as critical mediators of GLP1RA effects, providing key insights into the neural circuits underlying obesity therapeutics^15^.

Despite these advancements, GLP1RAs require continued administration, as discontinuation leads to weight regain^16^. Additionally, while patients experience substantial initial weight loss, this effect plateaus even with continued treatment^17^. Understanding the intracellular mechanisms that dictate how GLP-1 drugs work may offer avenues to address these issues. GLP1Rs are G-protein-coupled receptors (GPCRs) and previous studies largely in peripheral tissues report that GLP1R agonism increases levels of the secondary messenger cyclic adenosine monophosphate (cAMP) through the stimulatory alpha subunit G protein (Gs) protein signaling pathway to mediate its effects^18–22^. While some evidence suggests that GLP1RAs elevate cAMP in hindbrain neurons^18,23^, its role in mediating semaglutide’s effects on neuronal activity and weight regulation remains unclear. Given that cAMP can act on different timescales— modulating rapid neuronal excitability as well as longer-term cellular adaptations^24–28^— understanding the role of intracellular signaling mechanisms in regulating neuronal responses to semaglutide is crucial toward understanding the efficacy of GLP1-based drugs.

In this study, we employ two-photon imaging, viral genetic manipulations, and pharmacological approaches to systematically investigate the intracellular signaling mechanisms underlying how semaglutide drives weight loss through its primary site of action in the brain, the AP. Using anatomical mapping, fiber photometry and activity-based cell population capture, we identify the external lateral PBN (elPBN)—a satiety promoting hub—as a functional downstream correlate recruited by semaglutide-induced cAMP signaling in the AP. Our findings reveal the molecular and cellular mechanisms by which semaglutide acts through a cAMP-dependent hindbrain circuit to drive neuronal activation and weight loss, uncovering potential targets for enhancing obesity treatments.

## Results

### Local disruption of Gs signaling in the dorsal vagal complex impairs energy balance and semaglutide-induced weight loss

To test whether semaglutide recruits the Gs pathway in hindbrain neurons to drive weight loss, we delivered a virus encoding Cre recombinase and a red fluorescent protein (AAV1-hsyn-cre-P2A-tdTomato) to the dorsal vagal complex (DVC) of *Gnas* conditional knockout mice^29^ (Gnas^fl/fl^) to selectively knockout the gene encoding the Gs protein (Fig. 1 a-c). Control mice were injected with a control virus lacking Cre (AAV1-hsyn-mCherry) (Fig. 1c). Quantification of the spread of viral expression of Cre across the AP and NTS of the DVC yielded two groups of mice: 1) a DVC^ΔGnas^ group that had Gs signaling disrupted in both the AP and NTS and 2) an AP^Miss^ group that lacked viral expression in the AP, while containing viral expression in the NTS (Fig. 1c-d). To measure the effects of semaglutide, at 3 weeks after viral injection, mice were placed on a 60% high fat diet (HFD) for 3-5 weeks as a model for diet-induced obesity (Fig. 1e)^30^. There were no significant differences in bodyweight gained on HFD across groups (Fig. 1e). Mice were then given daily subcutaneous (SQ) injections of semaglutide (12 µg/kg) for 14 days and changes in bodyweight and food intake were recorded (Fig. 1f). Similar to previous reports^5^, control mice on average lost 9.3±0.65% bodyweight. In contrast, DVC^ΔGNAS^ mice did not lose weight in response to semaglutide treatment (−0.01±0.65%), showing that Gs signaling in the DVC is necessary for the weight loss effects of semaglutide. Interestingly, AP^Miss^ mice were responsive to semaglutide in a manner that was not significantly different from controls (−7.5%±1.4), suggesting that preservation of Gs signaling in the AP alone is sufficient to drive the full weight loss effects of semaglutide (Fig. 1f). In concordance with this, we observed that the degree of AP *Gnas* knockout, represented by percentage of viral expression in that region, is significantly correlated to weight change in response to semaglutide, whereby a higher degree of AP hit was predictive of less weight lost on semaglutide (Fig. 1g). NTS viral expression was not correlated with bodyweight responses to semaglutide but was significantly correlated with bodyweight changes to HFD (Extended Data Fig. 1a). Using a multiple linear regression model, we found that the percentage of AP *Gnas* knockout, but not NTS, is a significant predictor for change in bodyweight in response to semaglutide (Fig. 1h). Percentage of NTS *Gnas* knockout was instead a significant predictor for changes in bodyweight to HFD, suggesting Gs signaling in the NTS may play a role in bodyweight regulation (Extended Data Fig. 1b).

**Figure 1.**
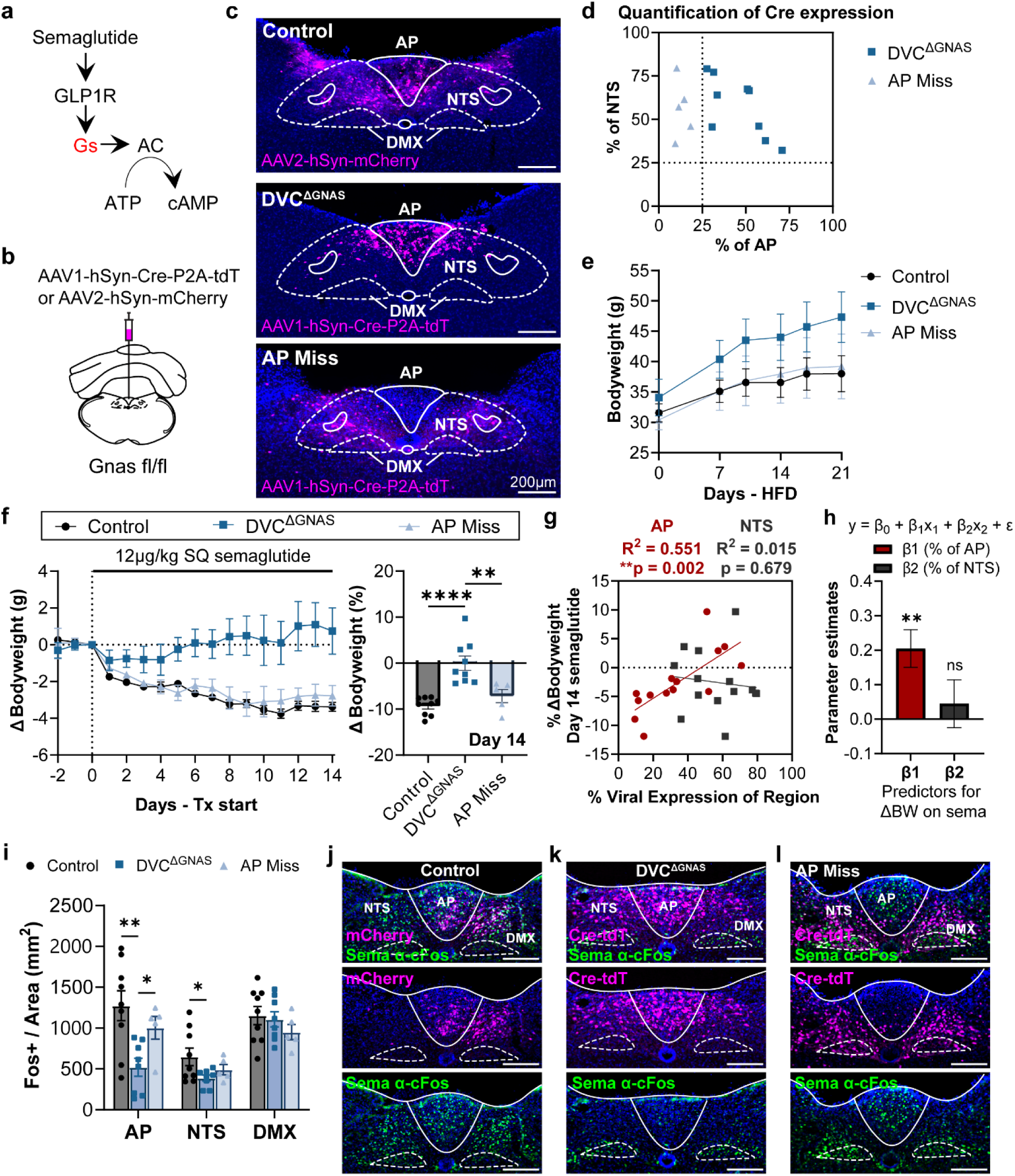
Local knockout of Gs signaling in the dorsal vagal complex attenuates semaglutide induced weight loss. **(a)** Schematic representation of the Gs-cAMP pathway. Semaglutide binds the GLP1R and triggers the stimulatory Gs protein to activate adenylyl cyclase (AC). Activated AC then converts adenosine triphosphate (ATP) to cAMP. **(b)** Virus and injection strategy for expressing Cre recombinase or mCherry control in the dorsal vagal complex of Gnas^fl/fl^ mice. **(c)** Representative images of viral expression in the three groups of mice: (top) Control (n=9 mice), (middle) DVC^ΔGNAS^ (n=9 mice), and (bottom) AP Miss (n=5 mice). Scale bar = 200 µm. **(d)** Quantification of extent (%) of Cre viral expression in area postrema (AP) and nucleus of the solitary tract (NTS) in DVC^ΔGNAS^ (square) and AP Miss (triangle) groups. **(e)** Bodyweight in grams (g) of the three groups of mice over three weeks on high fat diet (HFD) **(f)** Left: mean change in bodyweight (grams) from baseline on two week daily subcutaneous (SQ) injections of semaglutide (12 µg/kg). Right: bodyweight (%) changes on day 14 of semaglutide treatment. Points represent individual mice. Ordinary One-way ANOVA with Tukey’s multiple comparisons test (**p=0.0045, ****p<0.0001). **(g)** Simple linear regression of extent of viral expression in either AP (red) or NTS (gray) with the day 14 percent change in bodyweight to semaglutide treatment. AP: slope is significantly different from zero (**p=0.0024). NTS: slope is not significantly different from zero (p=0.6788). Slopes for AP and NTS are significantly different, p=0.0327. **(h)** Multiple linear regression for weight loss on semaglutide with predictors: β_1_ (% of AP viral expression) and β_2_ (% of NTS viral expression) (**p=0.0032, ns p=0.5353) **(i)** Quantification of semaglutide induced Fos in the DVC across AP, NTS, and the dorsal motor nucleus (DMX). Two-way repeated measures ANOVA test with Tukey’s multiple comparisons test (AP:**p=0.0038, *p=0.0241; NTS: *p=0.0443). **(j-l)** Representative images of Fos antibody staining in DVC quantified in **(i)** with the control **(j)**, DVC^ΔGNAS^ **(k)**, and AP Miss **(l)** groups. Top: images with 3 colors merged with DAPI in blue, middle: viral expression (magenta), bottom: Fos (green) antibody labeling to semaglutide injection. Scale bar = 200 µm. All data are represented as mean ± S.E.M. See also Extended Data Fig. 1.

To further assess the differences in these three groups of mice, we examined Fos protein expression—a proxy for neuronal activity—in response to SQ injections of semaglutide across the three subregions of the DVC (AP, NTS, and dorsal motor nucleus (DMX)) (Fig. 1i-l). Control mice exhibited high amounts of Fos in both the AP and NTS in response to semaglutide and knockout of Gs (DVC^ΔGNAS^ group) significantly attenuated Fos neuronal activation in both the AP and NTS (Fig. 1i-k). In agreement with the weight loss responses, Fos in the AP^Miss^ group was not significantly different from controls in both the AP and the NTS (Fig.1i, l). There were no significant differences in DMX Fos expression across the three groups (Fig. 1i-l). Taken together, these results support that semaglutide signals through the Gs pathway, particularly in the AP, to drive changes in Fos neuronal activation and weight loss in response to semaglutide. These results agree with previous reports that large molecular weight GLP1R agonists are unlikely to cross the blood-brain-barrier, and thus target and signal through circumventricular organs like the AP^5,31,32^.

### Semaglutide activates both Gs and Gq pathways to drive spike-dependent and spike-independent calcium increases in AP^Glp1r^ neurons

Given that semaglutide targets GLP1Rs, which are canonically thought to signal through the Gs pathway^33–35^, we next assessed whether Gs is necessary for semaglutide mediated neuronal activation in Glp1r-expressing cells. Fos activation to semaglutide was measured in mice lacking *Gnas* in all Glp1r-expressing cells (Glp1r-ires-cre: Gnas^fl/fl^) and compared to saline or semaglutide Fos responses in wildtype littermate controls (WT: Gnas^fl/fl^) (Extended Data Fig. 1c-f). Consistent with previous reports, in wildtype mice, semaglutide robustly activates AP, NTS, elPBN, CeA, and SFO compared to their saline injected counterparts (Extended Data Fig. 1c-d, and 1f). Global knockout of Gs in Glp1r-expressing cells abolished brain-wide semaglutide-induced Fos activation to levels similar to that of saline injected controls, suggesting that Gs signaling specifically in Glp1r-expressing cells is necessary for semaglutide mediated Fos neuronal activation (Extended Data Fig. 1e-f). Glp1r-ires-cre x Gnas^fl/fl^ mice exhibited hyperglycemia, likely due to the known role of Gs pathway in the pancreas for regulating blood glucose levels and thus were not used for behavioral studies (Extended Data Fig. 1g)^36,37^.

To further dissect how Gs is involved in semaglutide’s intracellular signaling pathway that gives rise to neuronal activation, we employed two-photon calcium imaging of *ex vivo* brain slices containing DVC from Glp1r-ires-cre: soma-targeted GCaMP8s (soma-GC8s) transgenic mice that had intact Gs signaling (GnasWT) or lack thereof (GnasKO) (Fig. 2a-c). Single cell calcium dynamics of Glp1r-expressing DVC (DVC^Glp1r^) cells were imaged during a 10 min wash-on of 100 nM semaglutide and cell responses were confirmed using a positive control wash-on of 10 mM KCl—which drives robust increases in fluorescence—at the end of each imaging session. All individual cell responses were first normalized to their max responses during the KCl wash-on period and then combined to perform unsupervised hierarchical cluster analysis. Clustering of 2,226 DVC^Glp1r^ cells (1,129 GnasWT cells and 1,007 GnasKO cells) yielded five clusters visualized by dendrogram and plotted as a heatmap and average response profile (Fig. 2d-e, Extended Data Fig. 2a-c). The cells from these clusters were then assigned as cell-type responders based on the area under the curve of their semaglutide response comprising: (1) low amplitude, (2) mid-amplitude, (3) high amplitude, (4) transient, and (5) inhibitory responders (Fig. 2f). Importantly, cells from both the AP and NTS were represented in all clusters and cells from all slices were represented in all clusters suggesting that the clusters were not skewed by batch effects (Extended Data Fig. 2e-f). To explore differences in the representation of GnasWT and GnasKO cells across clusters, we first compared the percentage of either GnasWT or GnasKO cells in each cluster by slice (Extended Data Fig. 2d). This revealed a higher percentage of GnasKO cells in the low-amplitude responder cluster 1 (c1), whereas there was a higher percentage of GnasWT cells found in the high-amplitude responder cluster 3 (c3). This trend was observed at the population level and percentage per slice (Extended Data Fig. 2d). Given our observation that Glp1r-ires-cre:Gnas^fl/fl^ mice that were injected with SQ semaglutide displayed no Fos neuronal activation (Extended Data Fig. 1f), we hypothesized that c1 low-amplitude responders may represent calcium responses that lack neuronal firing strong enough to drive Fos expression, whereas c3 high-amplitude responders may represent calcium responses that reflect increased neuronal firing in response to semaglutide.

**Figure 2.**
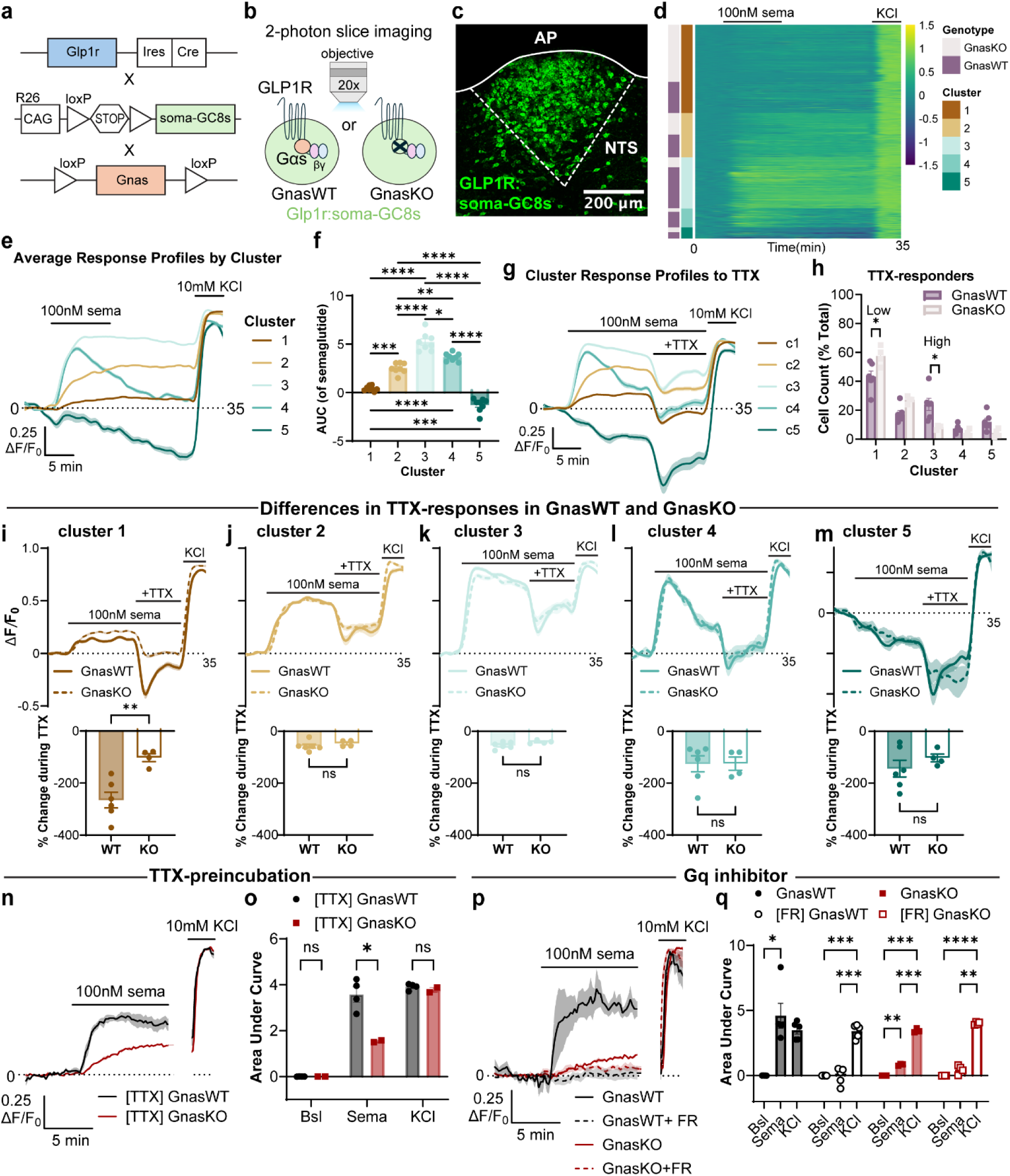
Semaglutide signals through Gs and Gq pathways to drive spike-dependent and spike-independent calcium dynamics in DVC^Glpr^ cells. **(a)** Schematic diagram of transgenic crosses used. Top: IRES-Cre fusion protein is driven under the direction of the *Glp1r* promoter, middle: CAG promoter and *loxP*-flanked STOP cassette upstream of the gene encoding soma-targeted GCaMP8s all inserted into the Rosa 26 (R26) locus, bottom: exon 1 of the *Gnas* gene is flanked by *loxP* sites. GnasWT mice are a cross of the first two transgenic lines and GnasKO mice are a cross of all three lines. **(b)** Schematic representation of the 2-photon slice imaging experimental paradigm of calcium transients in Glp1r-expressing DVC slices (Glp1r:soma-GC8s) with Gs intact (GnasWT) or absent (GnasKO). **(c)** Representative imaging field of view of the AP and NTS in a GnasWT slice. **(d)** Heatmap of calcium fluorescence of all 2,226 DVC^Glp1r^ cells (GnasWT: n=1,129 cells, 5 slices, 3 mice; GnasKO: n=1,007 cells, 3 slices, 3 mice) ordered by cluster (1-5) and genotype. **(e)** Average calcium response profile of cells by cluster. **(g)** Area under the curve quantification during the 10-minuted semaglutide (100 nM) wash per cluster per slice. Repeated measures one-way ANOVA test with Tukey’s multiple comparisons test (*p<0.05, **p<0.01, ***p<0.001, ***p<0.0001). **(g)** Average response profile by cluster of 2,292 DVC^Glp1r^ cells (GnasWT: n=1,154 cells, 6 slices, 3 mice; GnasKO: n=1,138 cells, 4 slices, 2 mice) based on their responses to semaglutide. **(h)** Percentage of cells in each cluster per slice by genotype. Two-way ANOVA with Benjamini, Krieger, and Yekuteli two-stage step-up method to correct for multiple comparisons (cluster 1: *p=0.0230, cluster 3: *p=0.0128). **(i-m)** Top: average response profile of each clusters response to tetrodotoxin (TTX; 500 nM) application separated by genotype. Bottom: quantification of change in fluorescence in response to TTX application per slice. Two-tailed unpaired t-test (cluster 1: **p=0.0034). **(n)** Mean calcium response in of semaglutide-responsive GnasWT (black) or GnasKO (red) cells following a 30-minute preincubation with TTX. **(o)** Area under the curve quantification per slice of bulk calcium fluorescence changes in the AP during the 5-minute baseline period (Bsl), the 10-minute semaglutide period (Sema) and the 5-minute KCl period (KCl) in GnasWT (black; n=4 slices, 2 mice) and GnasKO (red; 2 slices, 2 mice). Two-way ANOVA with Bonferroni’s correction (Sema: *p=0.0247). **(p)** Mean calcium response in of semaglutide-responsive GnasWT (black) or GnasKO (red) cells in ACSF (solid lines; GnasWT: n=5 slices, 3 mice; GnasKO: n=3 slices, 3 mice) or following a 45-minute preincubation with the Gq inhibitor FR (1 µM) (dotted lines; GnasWT: n=5 slices, 3 mice; GnasKO: n=4 slices, 2 mice). **(q)** Area under the curve quantification per slice of bulk calcium fluorescence changes in the AP from slices in ACSF (filled bars and points) or in the presence of FR (open bars and points). Bsl: 5-minute baseline period, Sema: 10-minute semaglutide period, KCl: 5-minute KCl period. Two-way ANOVA with Tukey’s multiple comparisons test (*p<0.05, **p<0.01, ***p<0.001, ****p<0.0001). All data are represented as mean ± S.E.M. See also Extended Data Fig. 2 and 3.

To test this, we again performed two-photon slice imaging of calcium responses of GnasWT and GnasKO cells during semaglutide wash-on. To assess the presence of spike-dependent responses to semaglutide, we subsequently bath applied the selective, high affinity voltage-gated sodium channel blocker tetrodotoxin (TTX; 500 nM) to block action potential firing and propagation (Fig. 2g). Hierarchical cluster analysis of 2,292 DVC^Glp1r^ cells (1,154 GnasWT cells and 1,138 GnasKO cells) based on their responses to semaglutide yielded the same five clusters observed previously (Fig. 2g). We observed a similar amount of TTX-responsive and TTX-nonresponsive cell responses regardless of genotype (Extended Data Fig. 2g). Following separation of TTX-responsive cells from each cluster for further analysis, we observed a significantly higher percentage of TTX responsive GnasWT cells in the high amplitude cluster (c3), supporting that GnasWT slices have an increased number of cells with neuronal firing in response to semaglutide (Fig. 2h). Unexpectedly, in the low amplitude cluster (c1), GnasKO slices had a significantly higher percentage of TTX responsive cells compared to GnasWT (Fig. 2h). To more carefully compare the differences in the TTX-responsive cells across GnasWT and GnasKO, clusters were separated by genotype (Fig. 2i-2m). This revealed that while GnasKO slices may have a higher percentage of TTX-responsive cells in cluster 1, the percent decrease in calcium signal in GnasKO cells during TTX application was significantly less than that of GnasWT cells (Fig. 2i). The larger decrease in signal in response to TTX in the GnasWT cells in c1 indicate that while c1 GnasWT cells have a low amplitude increase to semaglutide, they are spiking significantly more than GnasKO cells. This significant difference in degree of response to TTX was not observed in clusters 2-5 (Fig. 2j-m). Notably, out of total cells from each genotype, there was 48.9% of GnasWT cells and 51.7% of GnasKO cells that were unresponsive to TTX (Extended Data Fig. 2g). Calcium responses to semaglutide were still observed following a 30-minute pre-incubation with TTX, suggesting that there are both spike-dependent and spike-independent semaglutide-induced increases in calcium activity in GnasKO and GnasWT DVC^Glp1r^ cells (Fig. 2n, Extended Data Fig. 2h). However, the amplitude of spike-independent calcium release in GnasKO slices was significantly decreased compared to GnasWT slices, suggesting that the lack of Gs-mediated cAMP signaling blunts calcium responses to semaglutide (Fig. 2o).

The presence of semaglutide-responsive cells in Glp1r cells lacking Gs signaling was unexpected given that Gs is the known primary signaling pathway for GLP1Rs. However, some studies in peripheral cell culture suggest that GLP1Rs may couple to the G alpha q protein (Gq) under specific conditions, such as chronic hyperglycemia, thereby activating phospholipase C and resulting in intracellular calcium release from the endoplasmic reticulum^38,39^. Indeed, the remaining calcium transients observed in GnasKO DVC^Glp1r^ slices preincubated with TTX supports that calcium may be released through alternative pathways (Fig. 2n-o, Extended Data Fig. 2h). To investigate whether Gq signaling may play a role in driving calcium responses to semaglutide, GnasWT and GnasKO DVC slices were collected from Glp1r-ires-cre;GCaMP8s mice and pre-incubated with a selective Gq inhibitor (FR900359 or FR, 1 µM) (Fig. 2p-q, Extended Data Fig. 2i-j)^40^. Bulk calcium imaging of AP^Glp1r^ responses to semaglutide showed that pharmacological inhibition of Gq blocked calcium responses observed in both GnasKO and GnasWT slices (Fig. 2q). To test whether FR may have off-target effects on the Gs-cAMP pathway, a Cre-dependent virus carrying a fluorescent biosensor for cAMP (cADDis; AAV1-hSyn-flex-GreenDownwards-cADDis) was injected into the AP of Glp1r-ires-cre mice and cAMP responses to a 25-minute bath application of semaglutide (100 nM) were measured in the presence of FR (Extended Data Fig. 3a-b). The rapid adenylyl cyclase (AC) activator forskolin (Fsk, 10 µM) was bath applied at the end of each imaging session to drive cAMP production as a positive control for cell responses (Extended Data Fig. 4b). Importantly, while preincubation with FR could completely block calcium increases during a 10-minute bath application with semaglutide, FR did not block semaglutide-induced cAMP production, confirming that the Gs pathway was still intact (Extended Data Fig. 3c-g).These results suggest that while Gs is a necessary component to calcium and behavioral responses to semaglutide, Gq plays a significant role in the initial release of calcium in response to semaglutide.

Taken together, these data highlight a critical role for the Gs pathway for semaglutide-induced increases in neuronal firing and brain-wide Fos activation to semaglutide. They also reveal a previously unreported intracellular signaling mechanism utilized by semaglutide in the brain where spike-independent calcium is released through the Gq pathway.

### Semaglutide drives transient and sustained increases in cAMP and calcium levels in AP^Glp1r^ neurons

Following Gs-coupled GLP1R agonism, Gs binds to adenylyl cyclase (AC) which converts ATP to cAMP (Fig. 1a). In this signaling pathway, cAMP is an essential secondary messenger that recruits downstream biochemical cascades^33^. Given that Gs signaling is necessary in the AP for the weight-loss effects of semaglutide (Fig. 1), we next characterized cAMP responses to semaglutide in Glp1r-expressing AP neurons (AP^Glp1r^). A cre-dependent cADDis virus was delivered into the AP of Glp1r-ires-cre mice, and changes in cAMP fluorescence during a 25-minute wash-on of semaglutide (100 nM) in *ex vivo* brain slices were recorded using two-photon imaging (Fig. 3a-c). Single-cell cAMP dynamics revealed both transient cAMP responses (30.2% of 139 cells) that exhibited an initial increase followed by a return to baseline and sustained cAMP responses (69.8% of 139 cells) that showed an increase in cAMP signal which did not return to baseline levels in response to semaglutide (Fig. 3d-e, and g; see Methods). However, upon inspection of the cAMP response in individual cells, we noticed that there seemed to be heterogeneity within transient and sustained cAMP responses (Fig. 3d). To assess differences in how individual responses may change over time, the change from peak cAMP response (fraction of peak) of each cell was calculated at 5-minute increments during the semaglutide bath application (Fig. 3f). The cumulative distribution frequency of cAMP responses over time revealed that changes in cAMP levels were graded across the extent of drug wash, highlighting that transient and sustained categories are not binary but rather a gradient of individual cAMP responses to semaglutide (Fig. 3f).

**Figure 3.**
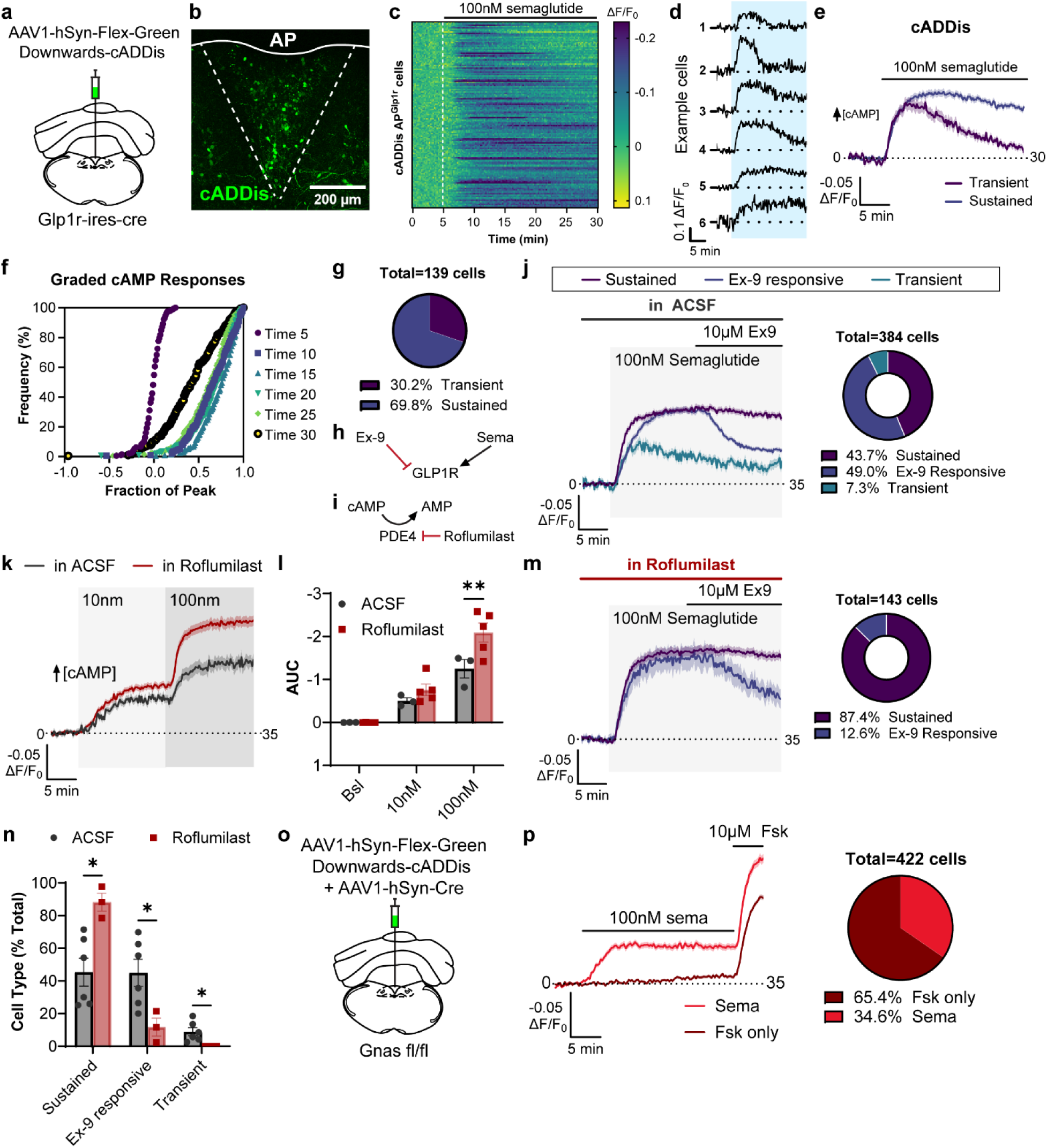
Semaglutide drives transient and sustained increases in cAMP in AP^Glp1r^ neurons. **(a)** Virus and injection strategy for expressing cADDis in Glp1r-expressing cells in the AP. **(b)** Representative image of cADDis expression (green) in the AP. Scale bar = 200 µm. **(c)** Heatmap of all cells (n= 139 cells, 4 slices, 2 mice) change in fluorescence in response to semaglutide (100 nM) bath application. The y-axis is inverted for clarity, as increases in cAMP are reflected as decreases in fluorescence in the GreenDownwards cADDis sensor. **(d)** Individual example traces from six cells in response to semaglutide wash (blue square). **(e)** Average response profile of transient cAMP responders (purple) and sustained cAMP responders (blue). **(f)** Cumulative frequency distribution of all the fraction of each cells peak response across 5-minute increments. **(g)** Pie chart of percentage of transient (purple) and sustained (blue) from total cells. **(h)** Schematic representation of the competitive antagonist exendin 9 (Ex-9) and the agonist semaglutide (Sema) for the GLP1R. **(i)** Left: average cADDis response profile of all cells (n=384 cells, 6 slices, 4 mice) response to semaglutide (100 nM) bath application followed by Ex-9 (10 µM) bath application in artificial cerebral spinal fluid (ACSF). Right: Pie chart of percentage of sustained (purple) and Ex-9 responsive (blue) and transient (teal) cAMP responders from total cells. **(j)** Schematic representation of the enzymatic breakdown of cAMP to AMP by phosphodiesterase 4 (PDE4). PDE4 activity is inhibited by Roflumilast. **(k)** Average cADDis response profile of all cells in response to 15-min bath application of 10 nM semaglutide followed by 15-min bath application of 100 nM semaglutide in ACSF (black line; n=45 cells, 3 slices, 3 mice) or in the presence of Roflumilast (10 µM; red line; n=107 cells, 5 slices, 4 mice). **(l)** Area under the curve (AUC) quantification per slice. Bsl: 5-minute baseline period, 10 nM: 15-minute 10 nM semaglutide period, 100nM: 15-min 100 nM semaglutide period. Two-way ANOVA with Bonferroni’s multiple comparisons test (**p=0.004). **(m)** Left: average cADDis response profile of all cells (n=143 cells, 3 slices, 3 mice) response to semaglutide (100 nM) bath application followed by Ex-9 (10 µM) bath application in the presence of Roflumilast (10 µM). Right: pie chart of percentage of sustained (purple) and Ex-9 responsive (blue) cAMP responders from total cells. **(n)** Percentage of cAMP type responses (sustained, ex-9 responsive or transient) per slice in ACSF (black) or Roflumilast (red). Two-way ANOVA with Bonferroni’s multiple comparisons test (Sustained: *p=0.0119, Ex-9 responsive: *p=0.0386, Transient: *p=0.0487). **(o)** Virus and injection strategy for expressing cADDis and Cre-recombinase in the AP of Gnas^fl/fl^ mice to measure cAMP responses in the absence of Gs. **(p)** Left: average response profiles of semaglutide-responders (Sema) or forskolin only (Fsk only) responders (n= 422 cells, 7 slices, 3 mice). Right: pie chart of percentage of Fsk only and Sema cAMP responders from total cells. All data are represented as mean ± S.E.M. See also Extended Data Fig. 4.

Previous reports in peripheral cells indicate transient cAMP responses could be a result of GLP1R desensitization from fast receptor internalization and degradation—a process that can, in part, depend on β-arrestins^41–43^. To investigate whether β-arrestins are necessary for the transient cAMP responses observed to semaglutide, we delivered a viral cocktail including Cre (AAV1-hSyn-Cre) and Cre-dependent cADDis to the AP of β-arrestin 1 and 2 conditional knockout mice (Barr1/Barr2^fl/fl^) (Extended Data Fig. 4a). Using two-photon imaging, we then recorded semaglutide-induced changes in cAMP fluorescence from cells lacking β-arrestin 1 and 2 (Extended Data Fig. 4c-d). Out of the 556 cells driven by the positive control Fsk, 11.9% had transient cAMP responses to semaglutide, 21% had sustained responses and 67.1% were responsive to forskolin, but not semaglutide (Fsk only) (Extended Data Fig. 4e). When comparing the proportion of transient responders out of total semaglutide responsive cells in slices from Barr1/Barr2^fl/fl^ mice to those from Glp1r-ires-cre mice, we observed no significant differences (Extended Data Fig. 4f), suggesting that transient cAMP responses to semaglutide may be regulated by β-arrestin independent processes.

One possible explanation for the sustained cAMP responses observed is that semaglutide remains bound to GLP1Rs located on the cell membrane. Another possibility is that semaglutide promotes prolonged cAMP signaling after GLP1R internalization, in which the GLP1R is recycled rather than degraded^42–46^. Thus, if semaglutide-induced sustained cAMP signaling requires continued receptor activation, we reasoned that bath application of the competitive GLP1R antagonist exendin-9 (Ex-9) should rapidly block the response (Fig. 3h). However, sustained cAMP signaling from GLP1R internalization should not be blocked by bath application of Ex-9 given that Ex-9 binds the extracellular domain of the GLP1R^47–49^. To test this, we imaged cADDis responses from AP^Glp1r^ neurons in *ex vivo* slices in response to a 15-minute bath application of semaglutide (100 nM) followed by the addition of Ex-9 (10 µM) for another 15 minutes (Fig. 3j). Out of the 384 cells recorded, 49% were Ex-9 responsive and rapidly inhibited in the presence of the antagonist, 43.7% continued to have a sustained cAMP response even in the presence of the antagonist, and 7.3% were transient cAMP responses, suggesting that membrane bound semaglutide represents only a portion of sustained cAMP responses (Fig. 3j). Notably, pre-incubation with the FDA-approved phosphodiesterase 4 (PDE4) inhibitor Roflumilast (10 µM)^50^— which prevents the breakdown of cAMP—not only abolished transient cAMP responses, but also increased the percentage of sustained responders and decreased the percentage of Ex-9 responsive cells across slices (Fig. 3i, 3m-n). The presence of roflumilast also significantly increased cAMP responses to semaglutide (100 nM) compared to control conditions, suggesting that cAMP levels are typically regulated by PDE4 activity (Fig. 3k-l).

Given our findings that semaglutide utilizes both Gs and Gq pathways in hindbrain neurons, we next probed whether Gs is necessary for the generation of semaglutide-induced increases in cAMP levels. To accomplish this, a viral mixture encoding Cre and a Cre-dependent cADDis was delivered into the AP of Gnas^fl/fl^ mice and fluorescent changes in cAMP levels to semaglutide were recorded using two-photon imaging (Fig. 3o). Unexpectedly, out of 422 cells that responded to Fsk, 34.6% still exhibited changes in cAMP fluorescence to semaglutide, suggesting that semaglutide can drive increases in cAMP levels independent of the Gs pathway (Fig. 3p). However, these cAMP responses were significantly decreased compared to those from AP^Glp1r^ slices with Gs intact, highlighting the importance of the Gs pathway in cAMP production (Extended Data Fig. 4g-i). Taken together, these results reveal the complex nature of cAMP production in AP^Glp1r^ cells in response to semaglutide, providing mechanistic insights into the development of future drug therapies. PDE4 inhibition can abolish transient responses and drive increased, more persistent cAMP levels to semaglutide. Moreover, semaglutide induces cAMP production possibly through Gs-dependent and Gs-independent pathways.

### cAMP elevation is necessary for semaglutide-induced sustained neuronal activity

cAMP plays a crucial role in facilitating diverse biological processes by acting as a bridge between external ligands and internal biochemical signaling transduction^24–28^. In this context, we next explored whether cAMP is necessary for the increases in neuronal activity driven by semaglutide. To inhibit cAMP levels, a Cre-dependent virus encoding a constitutively active PDE4 (PDE4-cat; AAV1-ef1α-DIO-mKate2-PDE4-cat)^51,52^ was delivered to the AP of Glp1r-ires-cre mice together with either Cre-dependent cADDis or a Cre-dependent calcium indicator GCaMP7s (AAV1-hsyn-Flex-GCaMP7s) to measure cAMP levels or neuronal activity in AP^Glp1r^ cells, respectively (Fig. 4a-b). Two photon imaging of cADDis fluorescence to a 10-minute bath application of semaglutide (1 µM) showed that PDE4-cat expression was sufficient to completely abolish cAMP responses in AP^Glp1r^ cells compared to slices where PDE4-cat was not present, confirming the efficacy of PDE4-cat to inhibit cAMP in this preparation (Fig. 4c-e). Subsequent two-photon imaging of calcium responses in AP^Glp1r^ cells lacking PDE4-cat expression (PDE4-) showed an expected increase in fluorescence to semaglutide, a response that was absent in AP^Glp1r^ cells expressing PDE4-cat (PDE4+) (Fig. 4f-j). Both cell types responded in a similar manner to bath application of KCl (10 mM) (Fig. 4h-i). These results show that cAMP elevation is a necessary step in driving increases in AP^Glp1r^ neuronal activity in response to semaglutide and that PDE4-cat is fully capable of abolishing these neuronal responses.

**Figure 4.**
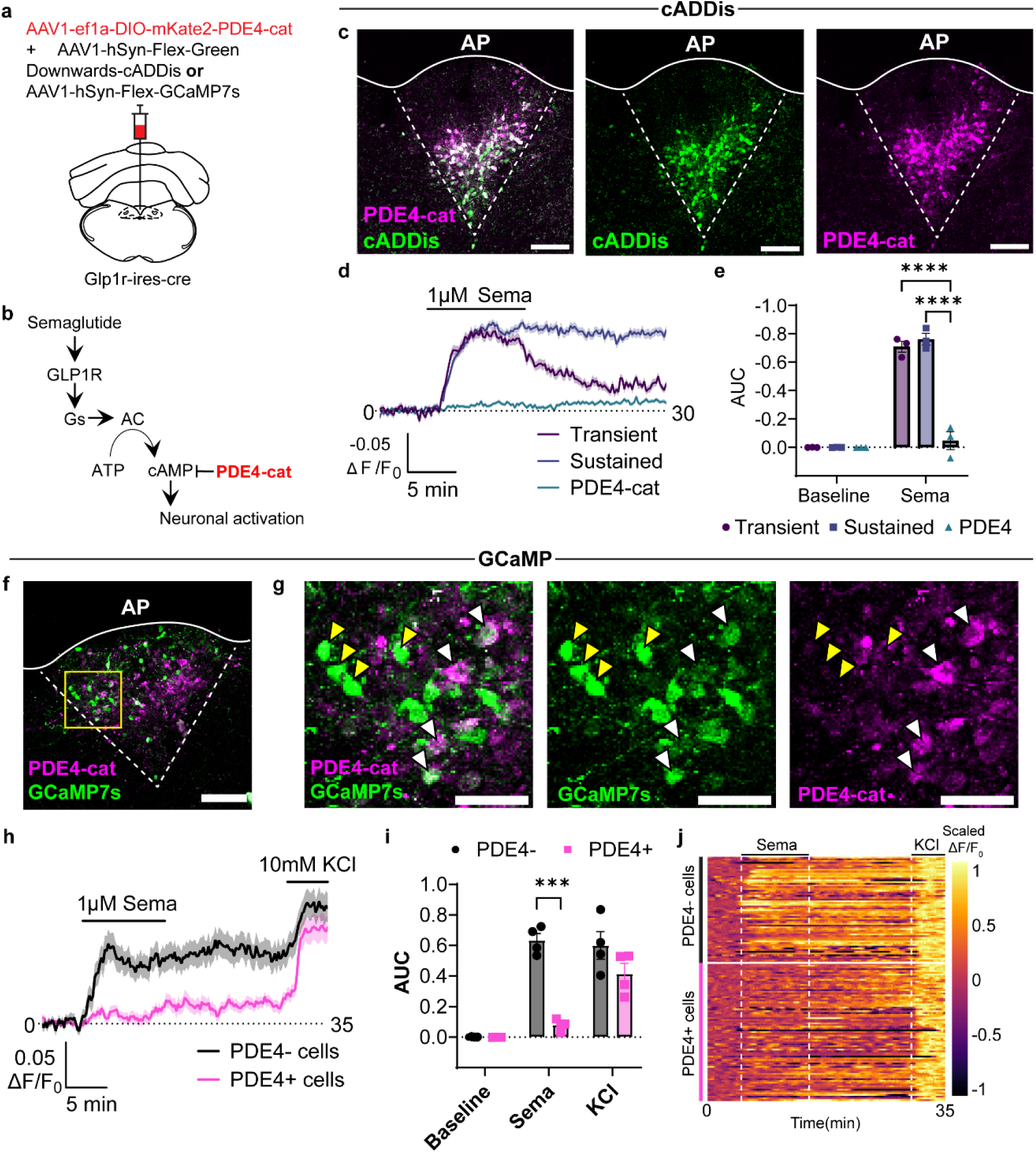
cAMP elevation is necessary for semaglutide-induced sustained neuronal activity. **(a)** Virus and injection strategy for expressing PDE4-cat with cADDis or GCaMP7s in Glp1r-expressing cells in the AP. **(b)** Schematic representation of the Gs-cAMP pathway. Semaglutide binds the GLP1R and triggers the stimulatory Gs protein to activate adenylyl cyclase (AC). Activated AC then converts adenosine triphosphate (ATP) to cAMP, which leads to downstream neuronal activation. cAMP is degraded by the constitutively active PDE4-cat. **(c)** Representative images of viral expression of PDE4-cat (magenta; right) and cADDis (green; middle) in AP^Glp1r^ cells. Scale bar = 200 µm. **(d)** Average response profiles of all cells cADDIs response to 10-minute bath application of semaglutide (1 µM) in cells lacking PDE4-cat expression (Transient: n=56 cells, 3 slices, 3 mice or Sustained: n=81 cells, 3 slices, 3 mice) or with PDE4-cat expression (n=140 cells, 3 slices, 3 mice). **(e)** Area under the curve (AUC) quantification per slice during 5-minute Baseline period and 10-minute Semaglutide (Sema) period. Two-way ANOVA with Tukey’s multiple comparisons test (****p<0.0001). **(f)** Representative image of viral expression of PDE4-cat (magenta) and GCaMP7s (green) in AP^Glp1r^ cells. Scale bar = 200 µm. Yellow box indicates field of view in (G). **(g)** Representative image of viral expression from yellow box in (F) showing PDE4-cat (magenta; right) colocalization with GCaMP7s (green; middle). White arrows indicate colocalization (PDE4+) and yellow arrows indicate GCaMP7s lacking colocalization (PDE4-). Scale bar = 100 µm. **(h)** Average response profiles of all cells GCaMP7s response to 10-minute bath application of semaglutide (1 µM) and 5-minute application of 10 mM KCl in cells lacking PDE4-cat expression (PDE4-: n=59 cells, 4 slices, 4 mice) or with PDE4-cat expression (PDE4+: n=102 cells, 4 slices, 4 mice). **(i)** Area under the curve (AUC) quantification per slice during 5-minute Baseline period , 10-minute Semaglutide (Sema) period and 5-minute 10 mM KCl period. Two-way ANOVA with Bonferroni’s multiple comparisons test (***p= 0.0009). **(j)** Heatmap of all GCaMP7s-expressing cells fluorescence response ordered by PDE4- (black) and PDE4+ (magenta). Dotted white lines indicate times of solution change. All data are represented as mean ± S.E.M.

### cAMP signaling in DVC^Glp1r^ neurons is necessary for semaglutide-induced weight loss and neuronal activation of downstream structures

To determine if cAMP is necessary for the weight-loss promoting effects of semaglutide, we again delivered Cre-dependent PDE4-cat to the DVC of Glp1r-ires-cre mice or that of wildtype littermates as control (Fig. 5a-b). Three weeks after viral injection, mice were fed 60% HFD for five weeks (Fig. 5c). Notably, PDE4 mice gained significantly more weight on HFD and exhibited increased food intake compared to their control counterparts (Fig. 5c). Both groups of mice were then treated with two weeks of daily SQ injections of semaglutide (12 µg/kg) or saline vehicle and bodyweight and food intake were tracked (Fig. 5d-e). Semaglutide treated control mice lost a significantly higher percentage of bodyweight compared to vehicle treated controls following two weeks of treatment (Fig. 5d). In contrast, PDE4 mice were completely unresponsive to semaglutide treatment and their change in bodyweight was not significantly different from saline injected controls (Fig. 5d). A similar trend in daily food intake was seen where control mice on semaglutide ate significantly less than saline treated controls, whereas PDE4 mice on semaglutide did not eat significantly less than their saline injected counterparts (Fig. 5e). Thus, cAMP signaling in DVC^Glp1r^ neurons is necessary for the weight loss effects of semaglutide.

**Figure 5.**
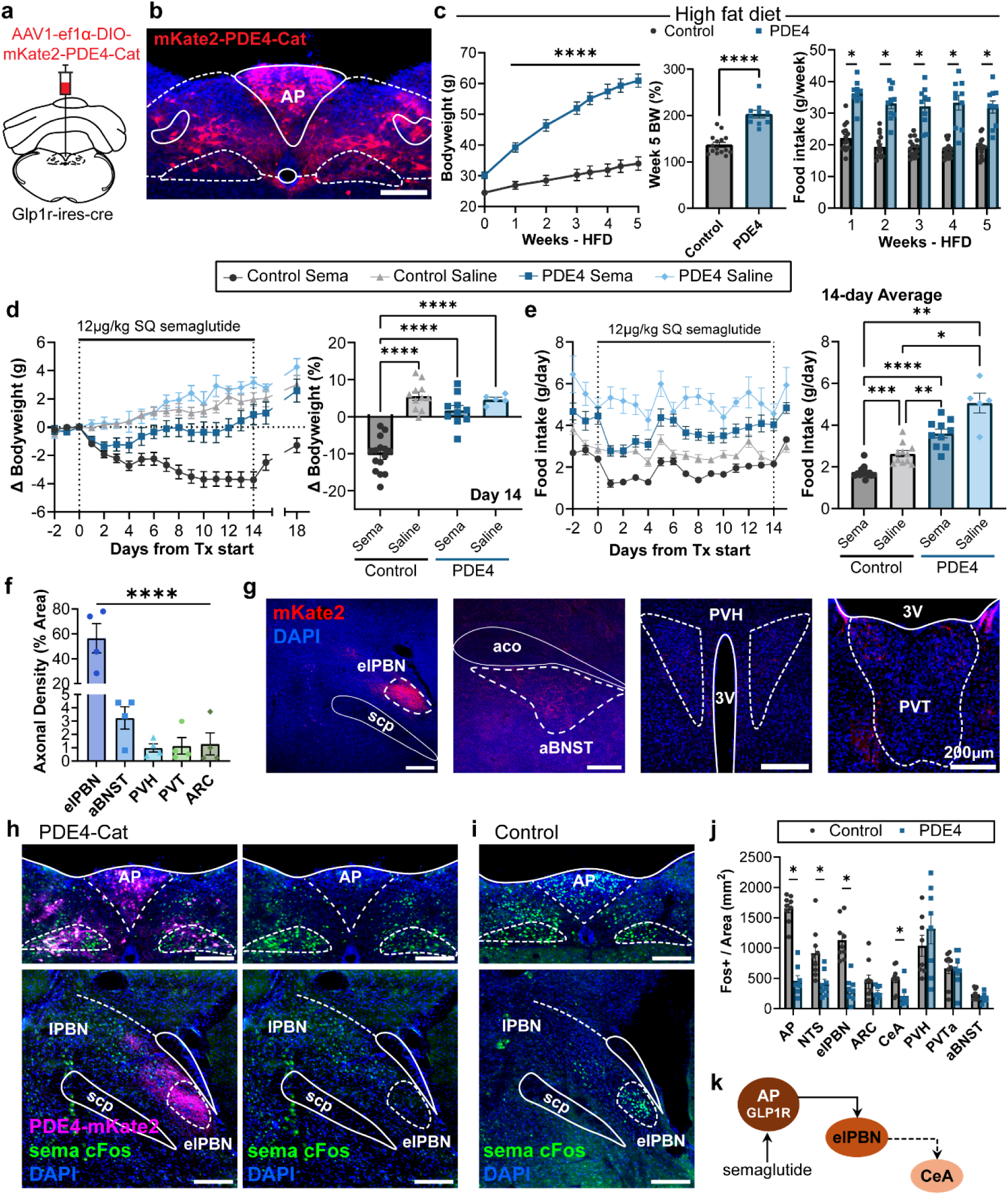
cAMP signaling in DVC^Glp1r^ neurons is necessary for semaglutide-induced weight loss and neuronal activation of downstream structures. **(a)** Virus and injection strategy for expressing PDE4-cat in the DVC of Glp1r-ires-cre mice. **(b)** Representative histological image of virus expression in Glp1r-expressing cells in the DVC with DAPI (blue) and PDE4-cat (red). Scale bar = 200 µm. **(c)** Changes in bodyweight in grams (left), week 5 percentage bodyweight (middle), and food intake (right) in the control (black; n=15 mice, 8 males) and PDE4 (blue; n=10 mice, 8 males) group on 5 weeks of high fat diet. Multiple unpaired t-tests with Welch’s correction (left: ****p<0.0001, right: *p<0.0001). **(d)** Left: change in bodyweight in the control group and PDE4 group in response to two weeks of daily SQ injection of semaglutide (12 µg/kg) or saline. Right: Day 14 change in bodyweight as percent. One-way ANOVA with Tukey’s multiple comparison’s test (****p<0.0001). **(e)** Left: Daily food intake in grams of the control group and PDE4 group in response to two weeks of daily SQ injection of semaglutide (12 µg/kg) or saline. Right: Mean food intake per day across 14 days of treatment. Brown-Forsythe and Welch’s one-way ANOVA with Dunnett’s T3 multiple comparisons test (*p= 0.0204, Control Sema vs PDE4 Saline:**p= 0.0097, PDE4 Sema vs Control Saline:**p= 0.0085, ***p= 0.0004, ****p<0.0001). **(f-g) (f)** Axonal density quantification of projections from AP^Glp1r^ neurons expressing mKate2-PDE4-cat in representative images found in **(g)**. Points represent individual mice. **(g)** elPBN: external lateral parabrachial nucleus; aBNST: anterior bed nucleus of the stria terminals; PVH: paraventricular hypothalamus; PVT: paraventricular thalamus; ARC: arcuate hypothalamus; scp: superior cerebellar peduncle; aco: anterior commissure; 3V: third ventricle. All scale bars = 200 µm. Repeated measures ANOVA with Tukey’s multiple comparisons test (elPBN vs. all other brain regions****p<0.0001). **(h-i)** Representative images of Fos immunostaining in response to semaglutide injections in mice expressing PDE4-cat in DVC^Glp1r^ neurons **(h)** or in Control mice **(i)**. (H) Left: PDE4-cat (magenta), semaglutide induced Fos (green), and DAPI (blue) merged. Right: Fos and DAPI of the same representative images from the right. All scale bars = 200 µm. **(j)** Quantification of brain-wide Fos expression in response to semaglutide in control (black) and PDE4 (blue) mice. Multiple unpaired t-test with Welch’s correction using the Holm-Šídák method (AP: p<0.000001, NTS: *p= 0.0379, elPBN: *p= 0.00016, CeA: *p=0.0379). **(k)** Schematic depiction that semaglutide binds and activates GLP1Rs in the AP. AP^Glp1r^ cells send downstream projections to recruit elPBN and indirectly recruit activation of the CeA. All data are represented as mean ± S.E.M.

To characterize the downstream projections of AP^Glp1r^ neurons, we performed serial sectioning from the brains of the PDE4 mice. Analysis of brain regions with expression of mKate2, a red fluorescent fluorophore that is tagged to PDE4, showed several labeled downstream structures including hypothalamic structures (arcuate (Arc) and the paraventricular hypothalamus (PVH)), paraventricular thalamus (PVT), and anterior bed nucleus of the stria terminalis (aBNST) (Fig. 5f-g). The region with the densest axonal projections was the external lateral parabrachial nucleus (elPBN), a region implicated in the control of food satiety and conditioned taste aversion (Fig. 5f-g)^10,14,53^. Next, to probe for functionally connected brain regions, we compared Fos responses to semaglutide in PDE4 mice and control mice across serial brain sections. Control mice showed high Fos expression to semaglutide in brain regions previously observed such as AP, NTS, elPBN, and CeA (Fig. 5i-j). Importantly, disruption of cAMP signaling in AP^Glp1r^ neurons in PDE4 mice significantly attenuated semaglutide Fos responses across all of these brain regions (Fig. 5h, j). The elPBN was the single region that exhibited both a robust innervation from AP^Glp1r^ cells and decreased Fos expression when cAMP was disrupted in AP^Glp1r^ cells, suggesting that the elPBN is a critical downstream structure recruited by semaglutide-induced cAMP signaling in AP^Glp1r^ neurons (Fig. 5k).

### elPBN^SemaTRAP^ neurons are rapidly recruited by peripheral semaglutide administration and are necessary for semaglutide induced conditioned taste aversion and weight loss

To probe the function of elPBN as a downstream node of semaglutide’s actions, we utilized the FosTRAP2 transgenic mouse model^54^ to gain genetic access to semaglutide-activated cells in the elPBN (Extended Data Fig. 5a). To validate this TRAP approach, we food deprived mice overnight to control for food intake across subjects and then performed SQ injections of saline vehicle (VehTRAP) for controls or semaglutide (SemaTRAP), followed by an intraperitoneal (IP) injection of 4-hydroxytamoxifen (4OHT) injection half an hour later to induce TRAPping. Six hours after 4OHT injections, we replaced the food in the cages. We first conducted a whole brain screening of neuronal activation to semaglutide by using FosTRAP2-Cre:Ai14 transgenic reporter mice to genetically label Fos activated cells with tdTomato. One week after performing TRAP, we injected SQ semaglutide into all mice, perfused the mice four hours later to perform Fos immunostaining, and identified brain regions differentially activated by semaglutide (Extended Data Fig. 5b). From this, we found the greatest differential colocalization in the AP, NTS, elPBN, CeA, and SFO supporting the specificity of this approach (Extended Data Fig. 5c-f).

To investigate whether SemaTRAPped cells are directly activated by semaglutide, we prepared SemaTRAP or VehTRAP brain slices of AP and elPBN from FosTRAP2.0-cre: soma-GCaMP8s transgenic mice and used *ex vivo* two-photon imaging to record calcium responses to semaglutide (Extended Data Fig. 5g-n). The percentage of semaglutide-responsive cells out of all KCl-responsive cells was significantly higher in AP^SemaTRAP^ slices compared to AP^VehTRAP^ slices, supporting our Fos results (Extended Data Fig. 5g-j). Importantly, elPBN^SemaTRAP^ slices had a similarly low percentage of semaglutide responders as elPBN^VehTRAP^ cells (Extended Data Fig. 5k-l), suggesting that elPBN^SemaTRAP^ cells are not directly responsive to semaglutide, but rather are recruited downstream of the primary site of semaglutide action—the AP.

Next, to better understand the time course of activity following peripheral semaglutide administration, we recorded *in vivo* neuronal responses from elPBN^SemaTRAP^ neurons using fiber photometry mediated calcium imaging by delivering a virus encoding a cre-dependent GCaMP7s to the elPBN and implanting a fiber above the injection site (Fig. 6a-b). Contrary to previous reports that central actions of GLP1RAs take hours or days^55–57^, elPBN^SemaTRAP^ neuronal activity increased within minutes following SQ semaglutide injection and stayed elevated for at least 20 minutes, suggesting that elPBN is a fast-acting neural substrate of semaglutide (Fig. 6c-d).

**Figure 6.**
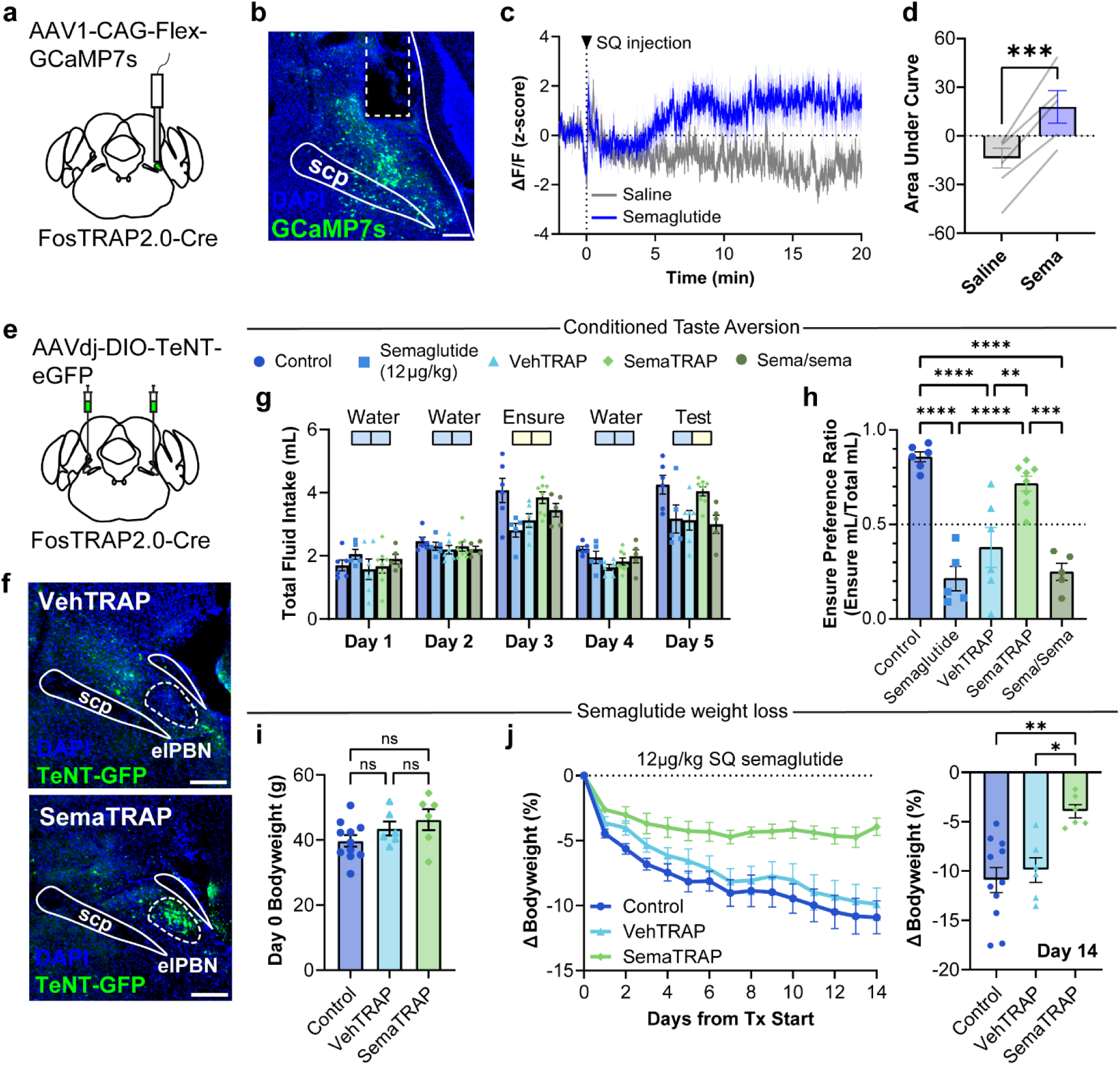
elPBN^SemaTRAP^ neurons are rapidly recruited by peripheral semaglutide administration and are necessary for semaglutide-induced conditioned taste aversion and weight loss. **(a)** Virus and fiber implant strategy for expressing GCaMP7s in the elPBN of FosTRAP2.0-cre mice to measure the activity of semaglutide-activated elPBN neurons. **(b)** Representative histological image of virus expression and fiber implant location above the elPBN with DAPI (blue) and GCaMP7s (green) from an elPBN^SemaTRAP^ mouse. Scale bar=200 µm. **(c)** Mean fiber photometry bulk calcium fluorescence (z-score) from elPBN^semaTRAP^ neurons (n=5 mice) in response to SQ saline (gray) or SQ semaglutide (blue; 120 µg/kg) injection. **(d)** Area under the curve quantification from time 0 to time 20 post injection. Gray lines represent individual mice. Two-tailed paired t-test (***p=0.001). **(e)** Virus and injection strategy for expressing TeNT bilaterally in the elPBN of FosTRAP2.0-cre mice. **(f)** Representative histological images of DAPI (blue) and TeNT viral expression (green) in the elPBN in VehTRAP (top) or SemaTRAP (bottom) mice. All scale bars = 200 µm. **(h-g)** Conditioned taste aversion behavioral paradigm with daily fluid intake (h) and ensure preference ratio (g) of five groups of mice. Control: Vehicle paired novel flavor group (n=6 mice); Semaglutide: Semaglutide (12 µg/kg) paired novel flavor group (n=5 mice); VehTRAP: Semaglutide (12 µg/kg) paired novel flavor with TeNT expressed in elPBN^VehTRAP^neurons (n=6 mice); SemaTRAP: Semaglutide (12 µg/kg) paired novel flavor with TeNT expressed in elPBN^SemaTRAP^ neurons (n=8 mice); Sema/sema: Semaglutide (12 µg/kg) paired novel flavor in mice with previous exposure to semaglutide (n=5 mice). One-way ANOVA with Tukey’s multiple comparisons test (**p= 0.0028, ***p= 0.0001; ****p<0.0001). **(i)** Bodyweight (grams) of mice on Day 0 of semaglutide treatment in (j). One-way ANOVA with Tukey’s multiple comparisons test (ns: not significant). **(j)** Left: mean weight responses to two weeks of daily SQ semaglutide (12 µg/kg) treatment in Control mice (n=11 mice), elPBN VehTRAP mice (n=6 mice) and elPBN SemaTRAP mice (n=6 mice). Right: change in bodyweight on Day 14 of semaglutide treatment. Points represent individual mice. One-way ANOVA with Tukey’s multiple comparisons test (*p=0.0191, **p=0.002). All data are represented as mean ± S.E.M. See also Extended Data Fig. 5.

In both humans and rodent models, use of GLP1R agonists is associated with aversive side effects, such as gastrointestinal disturbances and nausea or nausea-like symptoms^15,58,59^. Given that the elPBN has been implicated in both satiety and conditioned taste aversion, to determine whether elPBN^SemaTRAP^ neurons are required for semaglutide-mediated conditioned taste aversion, we chronically silenced these neurons by delivering a Cre-dependent virus expressing a tetanus toxin light chain construct (AAVdj-DIO-TeNT-eGFP) bilaterally to the elPBN and compared the responses of elPBN^SemaTRAP^ mice to either elPBN^VehTRAP^ mice or wildtype littermate controls (Fig. 6e-f). Using a 5-day conditioned taste aversion (CTA) protocol (see Methods), mice were first habituated to a two-bottle choice test using water. On day 3, mice were exposed to two bottles containing a highly palatable novel flavor ensure followed by a pairing with either a SQ saline injection for control mice or SQ semaglutide (12 µg/kg) for all other groups. On day 5, mice were presented with a two-bottle choice task with one bottle containing water and one bottle containing ensure and the ensure preference ratio was measured (Fig. 6g-h). Using this assay, semaglutide induced a robust conditioned taste aversion by decreasing the preference for the novel flavor when compared to control mice that instead developed a positive preference for the novel flavor (Fig. 6h). Chronic silencing of elPBN^SemaTRAP^ neurons blocked the development of CTA, an effect that was not seen upon chronic silencing of elPBN^VehTRAP^ neurons (Fig. 6h). Given that SemaTRAP mice are exposed to semaglutide for the first time during TRAPping (prior to the CTA paradigm), a fifth control group of wildtype mice (Sema/sema) was also tested in which they received a prior exposure to semaglutide in the same timeline as SemaTRAP mice (Fig. 6h). Importantly, Sema/sema mice still developed robust CTA that was not significantly different from the single exposure semaglutide group, confirming that elPBN^SemaTRAP^ neurons are necessary for the development of CTA to semaglutide (Fig. 6h).

To explore the role that elPBN plays in semaglutide-induced weight loss, the same groups of mice with chronic silencing of either elPBN SemaTRAP or VehTRAP cells and wildtype controls were treated with two weeks of daily SQ semaglutide injections. Control and VehTRAP mice lost 10.0±1.3% and 9.9±1.2%, respectively, whereas SemaTRAP mice were significantly less responsive to semaglutide losing 3.9±0.66% (Fig. 6i-j). Taken together, these results support that elPBN^SemaTRAP^ neurons are a functionally important downstream node from AP^Glp1r^ neurons and play an important role in mediating semaglutide induced conditioned taste aversion and its weight-loss effects. Moreover, these results highlight the functional specificity of the TRAP model for mapping, monitoring, and manipulating anti-obesity medication (AOM) sensitive neuronal populations.

## Discussion

In this study, we characterize the intracellular signaling mechanisms of semaglutide and demonstrate that it engages both Gs and Gq pathways in Glp1r-expressing AP neurons. While Gs signaling is the primary driver of cAMP production and is essential for the behavioral effects of semaglutide, we unexpectedly find that Gq signaling is required for the initial rise in intracellular calcium levels in response to the drug. Notably, cAMP serves as a central mediator linking these pathways to the recruitment of downstream brain regions, such as the elPBN, and is critical for the weight-loss effects of semaglutide. These findings provide mechanistic insight into the molecular, circuit-level, and behavioral effects of GLP1R-based anti-obesity therapeutics, offering a foundation for optimizing their efficacy and durability in future drug development.

### Gs signaling in the AP is necessary for semaglutide induced weight loss and Fos neuronal activation

A recent benchmark study identified DVC^Glp1r^ neurons as the essential neural population mediating the effects of GLP-1–based drugs^15^. Additionally, this study reported that AP^Glp1r^ and NTS^Glp1r^ neurons play distinct roles in mediating aversion and non-aversive satiety, respectively. In support of this, we find that Gs signaling in the DVC is required for semaglutide-induced weight loss. Notably, Gs signaling in the AP—but not the NTS—appears to be essential for these effects, as *Gnas* knockout in the AP significantly decreased semaglutide’s efficacy in reducing body weight. Conversely, preserving Gs signaling in the AP alone was sufficient for semaglutide to induce full weight loss effects and drive Fos neuronal activation in the hindbrain. These findings suggest that current large-peptide GLP-1 based drugs primarily signal through the AP and provide further evidence that these drugs do not cross the blood-brain-barrier to directly access non-aversive NTS^Glp1r^ neurons^5,60^.

Moreover, Gs knockout in NTS neurons caused increased body weight on a high-fat diet, suggesting that Gs signaling in the NTS is important for energy balance and body weight regulation. The NTS is well-positioned to regulate food intake and metabolic homeostasis, and our findings support models in which Gs signaling in the NTS acts as a negative regulator of energy balance, placing a brake on high fat diet intake^12,23,60–68^. These insights have implications for the development of small-molecule GLP-1 analogues, such as orforglipron^69^, which may have improved brain permeability compared to peptidergic GLP-1R agonists^70^. Our results dissecting how Gs signaling in distinct DVC subregions contributes to GLP-1–induced weight loss could inform strategies to enhance therapeutic efficacy while minimizing undesirable effects.

### Gs and Gq pathways contribute to semaglutide-induced calcium activity in the DVC

Semaglutide is well established to drive increases in Fos neuronal activation and recently has been shown to drive increases in calcium fluorescence *in vivo*^5,15,71^. However, the intracellular signaling mechanisms that give rise to semaglutide induced neuronal activation have remained largely unexplored. Previous studies in peripheral cells indicate that GLP1R can couple to multiple G-proteins, including Gs and Gq^38,39^—reports of Gi/o involvement are scarce^72^. Gs is the primary G-protein coupled to GLP1Rs, mediating the activation of adenylyl cyclase and subsequent production of cAMP^33,34^. Here, we show that Gs contributes to semaglutide-driven increased calcium fluorescence and neuronal firing, as GnasWT DVC slices exhibit greater responses compared to GnasKO slices. Additionally, we identify a robust role for Gq in semaglutide-driven calcium release in neurons, which may contribute to priming cells for action potential firing as release of intracellular stores of calcium through Gq signaling can lead to neuronal depolarization^25^. In the periphery, Gq-mediated GLP1R signaling is thought to occur primarily under conditions of ER stress or hyperglycemia, such as in diabetes^38^. In contrast to this, our findings demonstrate that Gq is a necessary component of GLP1R signaling in wild-type mice under baseline conditions. Future investigations should explore how the contributions of Gq in the brain may change across models for diabetes and obesity.

Notably, Gq-mediated calcium release appears to be involved in the rapid calcium response following semaglutide application, as preincubation with a Gq inhibitor could block calcium release in GnasWT slices during a 10-minute wash-on. Our interpretation of these results is that Gq may contribute to the initial depolarization of neurons through the calcium mobilization actions of inositol (1,4,5) triphosphate (IP_3_), working in tandem with the Gs pathway to drive increased neuronal firing in response to semaglutide^73^. Indeed, there is also previous evidence that protein kinase C (PKC), which is downstream of the Gq pathway, can directly activate AC providing a potential mechanism for synergistic cross-talk between the Gs and Gq pathways^74^. This function may be particularly relevant in the brain, where neurons—unlike non-excitable cells such as those in the skin or liver—use precise electrical activity to regulate function^75–77^. However, while we establish that Gs is necessary in the hindbrain for semaglutide-induced weight loss, the potential behavioral contributions of Gq signaling remain to be tested. Altogether, these findings highlight the interaction between Gs and Gq signaling in the brain^78^, providing mechanistic insights into how GLP1R agonists modulate neuronal circuits in a manner distinct from peripheral tissues.

### cAMP is necessary for semaglutide-induced weight loss and recruitment of a downstream neural network

A central finding of our study is that cAMP signaling is necessary for semaglutide-induced weight loss and downstream activation of other brain regions including the elPBN and CeA. Here, we provide the first systematic characterization of cAMP dynamics in intact brain tissue in response to semaglutide. Similar to peripheral tissues, we observe both transient and sustained cAMP responses^43,46,79^. PDE4 activity plays a crucial role in shaping these dynamics given that the PDE4 inhibitor roflumilast could abolish transient responses and enhance sustained cAMP elevations in response to semaglutide. Given that cAMP-biased agonism has emerged as a strategy to improve treatment efficacy of GLP-1 drugs^80–83^, our findings to drive more sustained cAMP dynamics may offer another avenue to further enhance weight loss. Interestingly, we find that β-arrestins did not play a discernable role in mediating transient responses in the brain. Given that PDE4 inhibitors such as roflumilast are FDA-approved for treatment of chronic obstructive pulmonary disease (COPD)^84,85^, their potential application in combination with Gs-coupled GPCR-based drugs—including GLP1R agonists—warrants further investigation.

We demonstrate that expression of a constitutively active PDE4 in DVC^Glp1r^ neurons not only abolishes neuronal activity, but also completely blocks the weight-loss effects of semaglutide. This complete suppression of calcium activity was not observed with Gs signaling knockout alone, suggesting that alternative pathways may contribute to cAMP production. Indeed, we found that semaglutide can still drive cAMP production in the absence of Gs, indicating a potential role for Gq-dependent regulation of cAMP generation^86,87^. Our results underscore that cAMP is a crucial second messenger for semaglutide to engage downstream brain regions and regulate body weight. These findings support current strategies to generate GLP-1 analogues through targeted modulation of cAMP dynamics^88–90^. Additionally, we present a model for studying cAMP dynamics in intact brain tissue, which could serve as a valuable tool for future *in vivo* investigations of cAMP signaling, particularly in the context of chronic GLP-1 treatment.

### TRAP-Based Targeting of Pharmacologically Activated Neurons and the Role of the elPBN

In this study, we introduce a TRAP-based approach to selectively target and investigate neurons activated by anti-obesity medications (AOM-TRAP), such as semaglutide, offering a valuable tool to explore the neural circuits underlying their effects. Using this method, we identify a population of neurons in the elPBN that mediate semaglutide-induced conditioned taste aversion and partially contribute to semaglutide’s weight-loss effects. This aligns with the well-established role of CGRP-expressing elPBN neurons in appetite regulation and aversion ^14,53^. Interestingly, CGRP-expressing neurons account for only ∼50% of elPBN neurons activated by the GLP1R agonist Exendin-4^14^, indicating that other neuronal subtypes contribute to the pharmacological effects. Our TRAP-based approach allows for the capture and transfection of a broader subset of neurons than those readily labeled by genetic markers such as *Cgrp*, enabling further investigation of their molecular identities and functional roles.

In addition to the elPBN and regions previously reported to bind GLP1RAs, including the Arc and PVH^5,55,91–96^, AOM-TRAP revealed other brain regions differentially activated by semaglutide that may contribute to the drug’s effects, including the central amygdala and subfornical organ. The CeA, implicated in emotional and reward-related feeding behaviors^97–99^, was recently linked to semaglutide-induced reductions in binge-like alcohol consumption in mice^100^, making it a compelling target for future studies on how GLP1R agonists influence food or drug motivation and intake. Similarly, the SFO, a key sensory circumventricular organ that also expresses GLP1Rs^101,102^, is involved in fluid balance and energy metabolism^103–107^, suggesting an additional pathway through which GLP1RAs may impact bodyweight.

### Limitations of the Study

Due to the technical challenges associated with imaging hindbrain and brainstem structures, our cAMP characterization was conducted in *ex vivo* brain slices. While this approach provides valuable insight, it may not fully capture the *in vivo* dynamics of cAMP signaling acutely and longitudinally across days or weeks. Recent studies have developed methods to overcome these technical barriers, and such techniques may be adapted in future work to investigate GLP-1 based cAMP dynamics *in vivo*^15,65,108^. Additionally, although GLP1R expression is highly conserved across species, there have been reports of species-specific differences that should be considered when interpreting our findings^109–114^.

### Future implications: Enhancing the efficacy of GLP1R-based therapies

Given that patients experience weight-loss plateaus and rebound weight gain upon discontinuation of GLP1RAs, our findings reveal potential avenues for improving drug efficacy. Developing pharmacotherapies that enhance both Gq and Gs signaling or inhibit PDE4 activity, selectively in the AP, to sustain cAMP signaling may optimize recruitment of neuronal activity to drive long-term weight loss promoting effects. Furthermore, interactions between combined pharmacotherapies that target multiple GPCRs on AP neurons may differentially influence cAMP and calcium dynamics, and our approaches are ideally suited to investigate these interactions. A deeper understanding of how GLP1R agonists activate intracellular signaling pathways in the brain represents a significant advancement for the development of next-generation therapeutics that offer enhanced efficacy and durability in the treatment of obesity.

## Methods

### Experimental Model and Subject Details

Mouse husbandry and experimental procedures were approved by the National Institutes of Health Animal Care and Use Committee. Mice were singly housed in 22-24°C with a 12-hour light/12-hour dark cycle. Mice were fed with *ad libitum* standard chow (Teklad F6 Rodent Diet 8664) and water, unless water-deprived for conditioned taste aversion experiments. For diet-induced obese mice, Rodent diet with 60 kcal% fat was used (Research Diets Inc. D12492). Mice used for surgery were at least 8 weeks old at the time of surgery. Mice of both sexes were 6-15 weeks old for behavioral experiments and 6-12 weeks old for two photon slice imaging experiments. Details on the number of mice used per experiment are found in the figure legends. Transgenic mouse lines used: *Rosa-Cag-LSL-soma-jGCaMP8s* (Ghitani et. al.); *Gnas^tm^*^5^*^.1Lsw^/J* (Lee Weinstein, JAX 035239); *Glp1r^tm^*^1^.^1^(cre)*^Lbrl^*/*RcngJ* (JAX 5776621); *βarr1/2-KO* (Robert Lefkowitz & Jurgen Wess, JAX 011130); *Fos^tm^*^21^(icre/ERT2)*^Luo^*/J (JAX 030323).

### Stereotaxic surgery

Surgeries were performed as previously described^115^. Briefly, mice were anesthetized with 2-5% isoflurane first in a chamber then mounted on a stereotaxic apparatus. A heating pad set at 37°C was placed below the mice. Mice were subcutaneously injected with 4 mg/kg meloxicam slow release and 1 mL 0.9% sterile saline. The skull was leveled and craniotomy was performed. Viruses were injected at a speed of 20 nL/min using a glass pipette with a tip diameter of 20–40 μm attached to an air pressure system, and the pipette was withdrawn 10 min after injection. The following coordinates relative to bregma were used: elPBN (AP -5.25, ML ±1.65, DV -3.8), or relative to obex^18,116^ were used: AP (AP 0.3, ML 0, DV -0.3). Mice were allowed to recover for at least 3 weeks post-surgery before further experiments. The following viruses were used for this study: AAV1-hSyn-Cre-P2A-dTomato (Addgene; Cat # 107738), AAV2-hSyn-mCherry (Addgene; Cat # 114472), AAV1-CAG-Flex-jGCaMP7s-WPRE (Addgene; Cat # 104495), AAV2/1-hSyn-DIO-GreenDownwardcADDis (Children’s Vector Core), AAV2/1-ef1a-DIO-mKate2-PDE4D3-Cat (Addgene; Cat # 169128), and AAVdj-CMV-DIO-eGFP-2A-TeNT (Wu Tsai Neurosciences Institute; Cat # GVVC-AA-71).

### Histology

Animals were deeply anesthetized with chloral hydrate (7% in saline) and transcardially perfused with phosphate-buffered saline (PBS; 1×, pH 7.4), followed by 10% neutral buffered formalin (NBF). After extraction, brains were post-fixed in 10% NBF at 4 °C for a minimum of 8 h and cryoprotected by transferring to a 30% PBS-buffered sucrose solution for a minimum of 24 h. Coronal brain sections (40-50 μm) were cut utilizing a freezing microtome (SM 2010R, Leica). For all Fos immunohistochemistry procedures, after extraction, brains were post-fixed in 10% NBF for roughly 2-4 h at room temperature (23-24 °C). Subsequently, brains were cryoprotected by transferring to 30% PBS-buffered sucrose solution for a minimum of 24 h. Using a freezing microtome (SM 2010R, Leica), coronal brain sections were cut into 40 μm sections and collected into four series.

### Immunohistochemistry

Sections were saturated with PBST (0.2% Triton X-100 in PBS) for membrane permeabilization for 30 min at room temperature. Brain sections were blocked in 10% normal goat serum (NGS) in PBST (0.1% Triton X-100 in PBS) for 1 h at room temperature, followed by incubation with a primary antibody in 10% NGS-PBST overnight at 4 °C. Sections were then washed with 0.1% PBST (4 x 10-15 min) and incubated with a secondary antibody at room temperature for 1 h. All incubations and washes were done over a shaker at low speed. After washing with PBS (4 x 10-15 min), sections were mounted onto glass slides, and a coverslip was added using VECTASHIELD® Vibrance™ Antifade Mounting Medium with DAPI (Vector Laboratories). Images were taken using a VS200 research slide scanner (Olympus). Image analysis and cell counting were performed using Qupath software (v0.5.1). The primary antibody used was anti-c-Fos (1:1,000 rabbit polyclonal, Millipore Sigma, ABE457). The fluorophore-conjugated secondary antibody (1:500, conjugated with goat anti-rabbit Alexa Fluor-488, A-11008) was purchased from Thermo Fisher Scientific. Antibodies were diluted in PBS with 10% NGS and PBST.

### Blood glucose measurements

Blood glucose measurements were taken at the end of the light cycle. Mice were gently restrained and the tail was sanitized with ethanol and a small portion of the tail tip was removed using scissors. The tail was gently massaged to generate a blood droplet and the blood sample was collected directly onto a glucose test strip (Contour). The test strip was inserted into a glucometer (Ascentia) and the blood glucose concentration was recorded.

### Two-photon imaging of *ex vivo* brain slices

As previously described^117^, animals were deeply anesthetized using isoflurane and rapidly decapitated. Brain tissue was extracted and immediately immersed in ice-cold, choline-based cutting solution (92 mM choline chloride, 10 mM HEPES, 2.5 mM KCl, 1.25 mM NaH2PO4, 30 mM NaHCO3, 25 mM glucose, 10 mM MgSO4, 0.5 mM CaCl2, 2 mM thiourea, 5 mM sodium ascorbate, 3 mM sodium pyruvate, pH 7.4). 200 µM thick sections were collected using a vibratome (Campden 7000smz-2) and then transferred to a warmed (36°C) choline cutting solution for 10-15 minutes, followed by continued recovery in 36°C oxygenated (95% O2/ 5% CO2) artificial cerebral spinal fluid (ACSF; 126 mM NaCl, 21.4 mM NaHCO3, 2.5 mM KCl, 1.2 mM NaHPO4, 1.2 mM MgSO4, 2.4 mM CaCl2, 10 mM Glucose) for 30 minutes. Slices were then kept at room temperature until use. To record, a single slice was transferred to a recording chamber (RC-26G; Warner Instruments) perfused with oxygenated ACSF at room temperature at a flow rate of approximately 2 mL/min. For imaging, a two-photon microscope (Olympus FVMPE-RS) equipped with an Insight laser was used at an excitation wavelength of 920 nm and resonant scanning was used to acquire 30 fps with a 20x 1.0 NA water-immersion objective (Olympus). The following drugs were used for this study: Semaglutide (Adipogen Life Siences; Cat # CAS0910463-68-2), Forskolin (Tocris; Cat # 1099), Tetrodotoxin citrate (Tocris; Cat # 1069), Roflumilast (Sigma Aldrich; Cat # 1099), and FR900359 (Cayman Chemical Company; Cat # 33666)

### Two-photon image processing & data analysis

Image registration for two-photon calcium images of brain slices was performed in FIJI (ImageJ). Max intensity projection of Z-stack images were first generated and XY rigid body registration was performed using the Turbo Registration plugin. Next, optical flow-based motion compensation correction was performed using the Flow registration plugin^118^. Cellpose^119^ was used to segment and identify cell bodies as regions of interest (ROIs), which were exported as masks to use in FIJI. The change in fluorescence was calculated by generating the average intensity projection of baseline florescence (F_0_) during the baseline period. Using the image calculator function in FIJI, F_0_ was subtracted from the fluorescent time course F and then the resulting image was divided by F_0_.

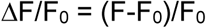

ROIs generated from Cellpose were then imported into the ROI manager and the mean intensity of each ROI was measured over time. Cells were designated as responsive if the change in fluorescence during the positive control (either KCl for GCaMP or Fsk for cADDis) was greater than ±2 standard deviations (S.D.) from the baseline period. Outside of hierarchical cluster analysis, cells were designated as responsive to semaglutide if the change from baseline during the first 10-minute wash on of the semaglutide was ±1 S.D. from the baseline period.

For transient and sustained cAMP responses, cells were defined as transient if the change in cADDis fluorescence returned to within 1 S.D. from the baseline period, whereas cells were defined as sustained if the change in cADDis fluorescence remained greater than 1 S.D. from the baseline.

For hierarchical cluster analysis of GnasWT and GnasKO cells, fluorescence values from each cell was first scaled to its maximum value during the KCl wash-on period. All values were then imported into R and traces were smoothed using a Savitzky-Golay filter (window=11, polynomial order=2). A distance matrix was computed containing the Euclidian distance between the rows of the fluorescence data and hierarchical clustering was performed using Ward’s method. Clustering was visualized using dendrogram, elbow plot of within-cluster-sum of squared errors, and silhouette plot and a cluster resolution of 5 clusters was selected. Data was further visualized by average cluster profile and by heatmap of all cells.

### Axonal density quantification

For axonal density quantification, images from serial sections of the whole brain were obtained using consistent exposure and scan settings and imported into FIJI (ImageJ). Images were thresholded using the Otsu’s method. Brain regions of interest containing projection fluorescence were measured using the measure function to calculate Area Fraction.

### Anti-obesity medication TRAP protocol

The day before TRAPping, 4-hydroxytamoxifen (4OHT, Sigma H6278) was prepared by following a previously described protocol^54,120^. Briefly, 4OHT was dissolved in 100% ethanol at a concentration of 20mg/mL at 37°C for 15 minutes. Corn oil was added to this solution and mixed at a concentration of 10 mg/mL and shaken with periodic heat for 15 minutes. The 4OHT solution was centrifuged briefly and placed on a 55°C heat block with the lid open until ethanol was fully evaporated. The final 10mg/mL 4OHT solution was placed in the 4°C fridge until use the following morning. For TRAP induction, mice were food restricted overnight 16-18 hours and then injected with subcutaneous saline or semaglutide. Thirty minutes post-injection, mice were injected intraperitonially with 4OHT at a dose of 50 mg/kg for Trap2.0-Fos-Cre:Ai14 or Trap2.0-Fos-Cre:soma-GCaMP8s mice and at a dose of 25 mg/kg for virally injected mice. Six hours after 4OHT injection, food was returned to the cage. Mice were given at least 7 days for recombination to occur prior to additional experiments.

### Fiber photometry

Mice were injected unilaterally into the elPBN (AP -5.25, ML -1.65, DV -3.8) with 150 nL virus (AAV1-CAG-). A 400 µm optical fiber was placed above the elPBN (DV -3.5) and secured with MetaBond (Parkell) and dental cement (Stoelting). Photometry recordings were performed Tucker-Davies Technologies system. Fluorescent signals were recorded using a commercial Mini Cube Fiber Photometry apparatus (Doric Lenses) connected to two continuously sinusoidally modulated LEDs (ThorLabs) tuned at 473 nm (211 Hz) and 405 nm (311 Hz), and two separate photoreceivers (2151, Newport Corporation). LEDs were coupled to a 400µm core, 0.48 NA optical fiber patch cord, which was mated to a matching brain implant in the mouse. GCaMP7s and autofluorescence (AF) signals were collected by the same fiber and focused onto separate photoreceivers. The data was analyzed by first applying a least-squares linear fit to the 405 nm signal to align it to the 470 nm signal. The resulting fitted 405 nm signal was then used to normalize the 473 nm as follows: ΔF/F = (473 nm signal − fitted 405 nm signal)/fitted 405 nm signal. Changes in fluorescence following behaviorally relevant events were quantified by measuring the area under the ΔF/F curve. All photometry data is presented as z-score of the ΔF/F from baseline (pre-stimulus) segments.

### Conditioned Taste Aversion Assay

Mice were water restricted daily for the 5-day duration of the assay outside of the duration of the 1-hour free access to fluid during the testing period, with *ad libitum* access to food. On days 1 and 2, during the 1-hour test period, mice were given free access to two bottles containing water to their home cage. On day three, mice were given 1-hour free access to both bottles containing the novel flavor (50% diluted ensure). Thirty minutes into the test period, mice were paired with subcutaneous injection of either vehicle control or semaglutide (12 µg/kg). On day 4, mice were again given water in both bottles during the test period. On day 5, mice were given a two-bottle choice assay during the test period, in which one bottle contained water and one bottle contained the 50% diluted ensure. Bottles were weighed before and after the testing period to obtain the daily fluid intake. Spouts were washed thoroughly and daily with Alconox and ethanol. Ensure preference ratio was calculated by the amount of ensure consumed over the total volume consumed from both bottles.

## Acknowledgements.

We thank Dr. Lee Weinstein for sharing the Gnas^fl/fl^ mouse line and Drs. Jurgen Wess and Robert J. Lefkowitz for sharing the Barr1/Barr2^fl/fl^ mouse line. We thank the support of the National Institute of Diabetes and Digestive and Kidney Disorders (NIDDK) animal facility staff. C.G. was supported by the National Institute of General Medical Sciences Postdoctoral Research Associate Training Program (Fi2-GM154675). The work from this study was supported by the NIDDK (Grants 1ZIA-DK075090-11 & 1ZIA-DK075169-2).

## Author contributions

C.G., A.L., and M.J.K. conceptualized the study. CG. and I.G. performed all experiments and histological analysis. C.G., C.L., and K.M. performed validation experiments. C.G. and I.G. performed all formal analysis and data visualization. A.L. and M.J.K. contributed resources used for this study. C.G., A.L., and M.J.K. wrote the paper. M.L.R. revised and edited the paper. M.L.R., A.L., and MJ.K. supervised the study.

## Additional information

Requests for further information and resources should be directed to and will be fulfilled by Dr. Michael J. Krashes. Reprints and permissions information is available at www.nature.com/reprints.

## Competing interests

The authors have no competing interests to declare.

**Statistical information.** All statistical analyses were imported and performed in GraphPad Prism. Statistical details and sample sizes are found in the figure legends. Normality tests were first performed to determine the appropriate statistical test to use. All data are presented as mean ± S.E.M. No assumptions or corrections were made prior to data analysis. All viral expression and optic fiber implant placements were verified by *post hoc* histology. Mice with incorrect virus expression location or optic fiber placement were excluded from the study. All experiments were replicated at least three times, and all subjects were randomly allocated to the different experimental conditions. The sample sizes used in our study were based on those in previously published studies or are about the same or exceed those estimated by power analysis (power = 0.8, α = 0.05). Significance levels are indicated as follows unless otherwise stated in the legends: *p<0.05, **p<0.01, ***p<0.001, ****p<0.0001.

**Extended data.** Extended data figures 1-5 and figure legends.

**Data availability.** All data is available upon request from corresponding authors.

